# An optimized pipeline for parallel image-based quantification of gene expression and genotyping after *in situ* hybridization

**DOI:** 10.1101/149591

**Authors:** Tomasz Dobrzycki, Monika Krecsmarik, Florian Bonkhofer, Roger Patient, Rui Monteiro

**Affiliations:** MRC Molecular Haematology Unit, MRC Weatherall Institute of Molecular Medicine, John Radcliffe Hospital, University of Oxford, Oxford, OX3 9DS, United Kingdom; BHF Centre of Research Excellence, Oxford, United Kingdom

**Keywords:** zebrafish, mutant, bias, genotyping, image quantification

## Abstract

Advances in genome engineering have resulted in the generation of numerous zebrafish mutant lines. A commonly used method to assess gene expression in the mutants is in situ hybridization (ISH). Because the embryos can be distinguished by genotype after ISH, comparing gene expression between wild type and mutant siblings can be done blinded and in parallel. Such experimental design reduces the technical variation between samples and minimises the risk of bias. This approach, however, requires an efficient method of genomic DNA extraction from post-ISH fixed zebrafish samples to ascribe phenotype to genotype. Here we describe a method to obtain PCR-quality DNA from 95-100% of zebrafish embryos, suitable for genotyping after ISH. In addition, we provide an image analysis protocol for quantifying gene expression of ISH-probed embryos, adaptable for the analysis of different expression patterns. Finally, we show that intensity-based image analysis enables accurate representation of the variability of gene expression detected by ISH and that it can complement quantitative methods like qRT-PCR. By combining genotyping after ISH and computer-based image analysis, we have established a high-confidence, unbiased methodology to assign gene expression levels to specific genotypes, and applied it to the analysis of molecular phenotypes of newly generated *lmo4a* mutants.

**SUMMARY STATEMENT:** Our optimized protocol to genotype zebrafish mutant embryos after in situ hybridization and digitally quantify the in situ signal will help to standardize existing experimental designs and methods of analysis.

## INTRODUCTION

The emergence of genome engineering technologies, including zinc-finger nucleases (ZFNs)(Doyon et al., 2008; Meng et al., 2008), transcription activator-like effector nucleases (TALENs)(Huang et al., 2011) and clustered regularly interspaced short palindromic repeats (CRISPR)/Cas9 (Hwang et al., 2013), has enabled zebrafish researchers to generate a wide range of mutant lines by precisely targeting genomic loci with high efficiency (Varshney et al., 2015). These methods provide alternatives to morpholino oligonucleotides (MOs), which have been used for gene knock down studies for over 15 years (Nasevicius and Ekker, 2000). In the light of recent concerns regarding the reliability of phenotypes induced by MOs (Kok et al., 2015), targeted gene knockouts have become a powerful tool to verify the transient MO phenotypes (Rossi et al., 2015) or to study genetic mutants, using MOs as a secondary tool (Gore et al., 2016). Low-cost protocols for DNA isolation (Meeker et al., 2007) and for genotyping of zebrafish embryos (Wilkinson et al., 2013) have allowed faster generation and analysis of new mutant lines.

One of the most widely used methods to analyse molecular phenotypes during embryonic development is *in situ* hybridization (ISH). This technique is used to detect spatial expression patterns and tissue-specific changes in mRNA levels. Relevant protocols with several extensions have been standardised (Westerfield, 2007) and numerous validated probes are curated on the ZFIN database (Howe et al., 2013). Using ISH to analyse MO phenotypes requires processing of the MO-treated and control samples separately, which may result in an inherent bias and an increased risk of technical variation. Lack of reported measures to reduce the risk of bias has recently been exposed in a meta-analysis of *in vivo* animal studies (Macleod et al., 2015). Genomic mutants offer a way to overcome these issues. Wild type and mutant embryos can be processed in one sample, assessed phenotypically in a blinded manner and then distinguished based on their genotype. This way, the technical variation between samples is minimised and phenotypic assessment is largely bias-free. A few publications have reported approaches to genotyping mutant lines after ISH, using either proteinase K treatment (Gore et al., 2016; Zhu et al., 2011) or commercial kits (Bresciani et al., 2014) for subsequent DNA extraction. However, the efficiency of these DNA extraction and genotyping methods has not been adequately demonstrated.

Reporting of phenotypes has usually been confined to one representative ISH image per condition, limiting the ability to quantitatively represent the variability of the phenotype. Approaches to score the expression levels as ‘high’, ‘medium’ or ‘low’ by eye (Blaser et al., 2017; Gao et al., 2016; Genthe and Clements, 2017; Peterkin et al., 2007; Place and Smith, 2017) are subjective and limited in how accurately they represent the effect of a knockout or a knockdown. Furthermore, visual scoring is inherently prone to poor reproducibility and low sensitivity. A more accurate way to quantitatively evaluate the change in gene expression levels involves counting the cells that contain the ISH signal (Espín-Palazón et al., 2014). However, this method is difficult to apply to compact anatomical structures and cell boundaries are hard to distinguish after ISH. Intensity-based image analysis using a selected region of interest (ROI) provides an objective alternative to visual scoring. Fan and colleagues have recently described a method using the ImageJ software to quantify ISH signal intensity in mouse embryos, where measurements were taken along a straight line drawn across the developing forelimb (Fan et al., 2015). Wen et al. have adapted this technique for use in the zebrafish to quantify gene expression levels in the mesoderm during early embryonic development (Wen et al., 2017). This approach can be further optimised to quantify gene expression after ISH in other regions and at other stages of the developing embryo.

Here we provide an optimised protocol for fast, inexpensive and highly efficient isolation of DNA from fixed zebrafish embryos for PCR and genotyping after ISH. This approach, based on the Hot Sodium Hydroxide and Tris (HotSHOT) method (Truett et al., 2000), is extremely reliable and allowed successful genotyping in 95-100% of the embryos. In addition, we propose a detailed step-by-step guide for gene expression intensity measurements based on modifications to the previously described quantitation methods (Fan et al., 2015; Wen et al., 2017). This pipeline (Fig. 1A) provides a tool for high-confidence, bias-free reporting of molecular phenotypes using standard ISH. We demonstrate its usefulness by assessing expression changes in *gpr65* morphants, and in *runx1* and *lmo4a* mutants.

**Figure 1.**
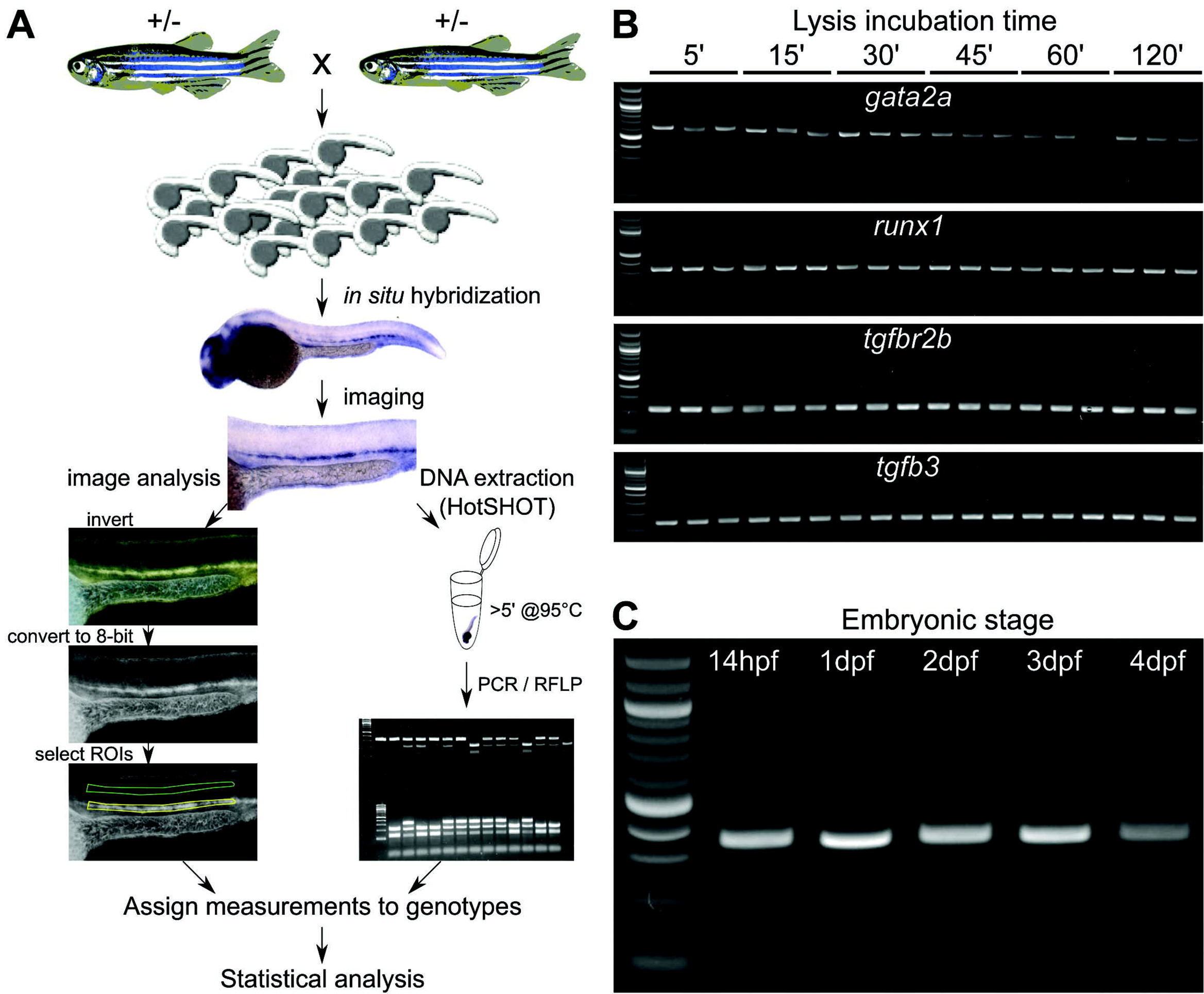
Overview of the method to extract DNA for genotyping zebrafish mutants after ISH and measure the mRNA levels. A) Embryos collected from an incross of fish heterozygous for a mutant allele are probed for the measured gene with a standard ISH protocol. After imaging in 100% glycerol, genomic DNA is extracted using the HotSHOT protocol by adding the lysis buffer directly to the embryo in a 0.2ml PCR tube, followed by a ≥5 min incubation at 95°C. This DNA is used for genotyping of the embryos by PCR and restriction fragment length polymorphism (RFLP). In parallel, the images for each embryo are inverted and converted to 8-bit greyscale. ROIs of identical shape and size containing the ISH signal (yellow) and background (green) are manually selected and measured. The measurements, assigned to corresponding genotypes, are statistically analysed. B) DNA fragments amplified from single fixed and ISH-probed embryos after incubation in HotSHOT lysis buffer for 5, 15, 30, 45, 60 and 120 min followed by JumpStart^™^ REDTaq^®^ ReadyMix^™^ PCR. Primer pairs designed for the *gata2a, runxl, tgfbr2b* and *tgfb3* produced 600, 300, 176 and 144 bp fragments, respectively. Three wild type embryos were used for each time point. First lane from the left: 100bp DNA ladder. C) DNA fragments amplified from single fixed and ISH-probed wild-type embryos after DNA extraction in HotSHOT lysis buffer for 5 min using a primer pair to amplify a 300bp fragment of the *runx1* locus with JumpStart^™^ REDTaq^®^ ReadyMix^™^ PCR. First lane from the left: 100bp DNA ladder. The lanes after the ladder represent embryos that are 14h, 1, 2, 3 and 4 days old and have been stored in glycerol at room temperature for 2 years.

## RESULTS

### HotSHOT genomic DNA extraction is suitable for PCR amplification in ISH-probed embryos

Our aim was to establish a fast and cost-effective way to extract DNA from individual embryos after ISH. First, we optimized the published HotSHOT protocol (Truett et al., 2000) for raw genomic DNA extraction, followed by JumpStart^™^ REDTaq^®^ PCR. By testing amplicons designed for four different loci, we found that a 5-minute incubation time in the lysis buffer was sufficient to extract DNA to amplify fragments ranging from 144 to 600 base pairs (bp) by PCR (Fig. 1B). Increasing incubation time did not increase DNA yield (Fig. 1B). Next, we tested the 5-minute DNA extraction method on embryos at different stages by amplifying a 300bp fragment from the *runx1* locus. We successfully extracted DNA from fish ranging from 14hpf to 5dpf that had been stored for up to 2 years in glycerol at room temperature (Fig. 1C; Table 1). Incubation for up to 30 min was sufficient to successfully genotype >96% embryos and incubation times longer than 65 min did not increase this efficiency (Table 1). When comparing two different commercially available PCR master mixes, we found that the JumpStart^™^ REDTaq^®^ ReadyMix^™^ performed more consistently on fixed ISH-probed genomic DNA samples than Phire^™^ Green HotStart II PCR Master Mix (Fig. S1). This observation might be due to differences in the optimal salt concentrations for the enzymes in each PCR mix.

**Table 1.** The efficiency of the HotSHOT protocol for post-ISH DNA extraction from zebrafish embryos at different stages depending on the lysis incubation time.

| Age \ Incubation time | ≤30 min | ≥65 min |
| --- | --- | --- |
| 22-26hpf | 99.3% (n=34-83; N=9) | 96.4% (n=56; N=1) |
| 28-30hpf | 99.3% (n=36-83; N=7) | 94.2% (n=59-67; N=8) |
| 32-48hpf | 100% (n=46-67; N=6) | 100% (n=38-67; N=2) |
| 4-5dpf | 96.6% (n=35-58; N=5) | 88.7% (n=55-61; N=2) |

n: number of embryos per experiment; N: number of independent experiments

### Digital quantification of the ISH signal in the dorsal aorta shows a significant decrease of dnmt3bb.1 mRNA levels in runx1^W84X/W84X^ mutants and a significant increase of runx1 expression in gpr65 morphants

To demonstrate the usefulness of our genotyping protocol in a known mutant, we imaged 130 embryos from the incross of *runx1^+/W84X^* heterozygotes (Jin et al., 2009), fixed at 33hpf and probed for *dnmt3bb.1* mRNA, a known downstream target of *runx1* within the haemogenic endothelium (Gore et al., 2016) (Fig. 2A). Scoring of the images as ‘high’, ‘medium’ or ‘low’ showed a Mendelian 1:2:1 distribution of phenotypes (Fig. S2A). We then genotyped all the imaged embryos with 100% efficiency using the above protocol and Restriction Fragment Length Polymorphism (RFLP)Fig. 2B). The observed Mendelian distribution of phenotypes, resulting from the first phenotypic assessment, did not entirely correspond to the respective genotypes. While the ‘low’ phenotype was significantly overrepresented in the homozygous mutant group (*X*^2^=95.3, d.f.=4; p<0.001), there was no significant difference in the distribution of ‘high’- and ‘medium’ -expressing embryos among wild type and heterozygous fish (*X*^2^=1.35, d.f.=1; p>0.2) (Fig. S2B). To implement a more quantitative assessment, we used Fiji (Schindelin et al., 2012) to digitally quantify pixel intensities of each embryo in the dorsal aorta region along the yolk sac extension prior to genotyping. This approach relied on allocating two separate ROIs to each image: one containing the staining (yellow ROI, Fig. 1A) and another containing an equal area of the embryo without any staining (green ROI, Fig. 1A). By subtracting the background value from the staining value, a number was assigned to each embryo. After genotyping, we compared the signal intensity values in wild type, heterozygous and mutant embryos. As expected, the *runx1* mutant embryos showed a statistically significant reduction by approximately 50% of *dnmt3bb.1* signal compared to wild types or heterozygotes (μ_wt_=54, μ_het_=50.1, μ_mut_=26.3; *F*=132.97, d.f.=2, 69.2; *p*<0.001)(Fig 2C). In contrast, there was no significant difference in signal intensity between wild types and heterozygotes (μ_wt_=54, μ_het_=50.1; *p*>0.3) (Fig. 2C). We found that the signal intensity values in all groups were very dispersed, with high coefficients of variation (24%, 22% and 21% for wild type, heterozygote and mutant groups, respectively).

**Figure 2.**
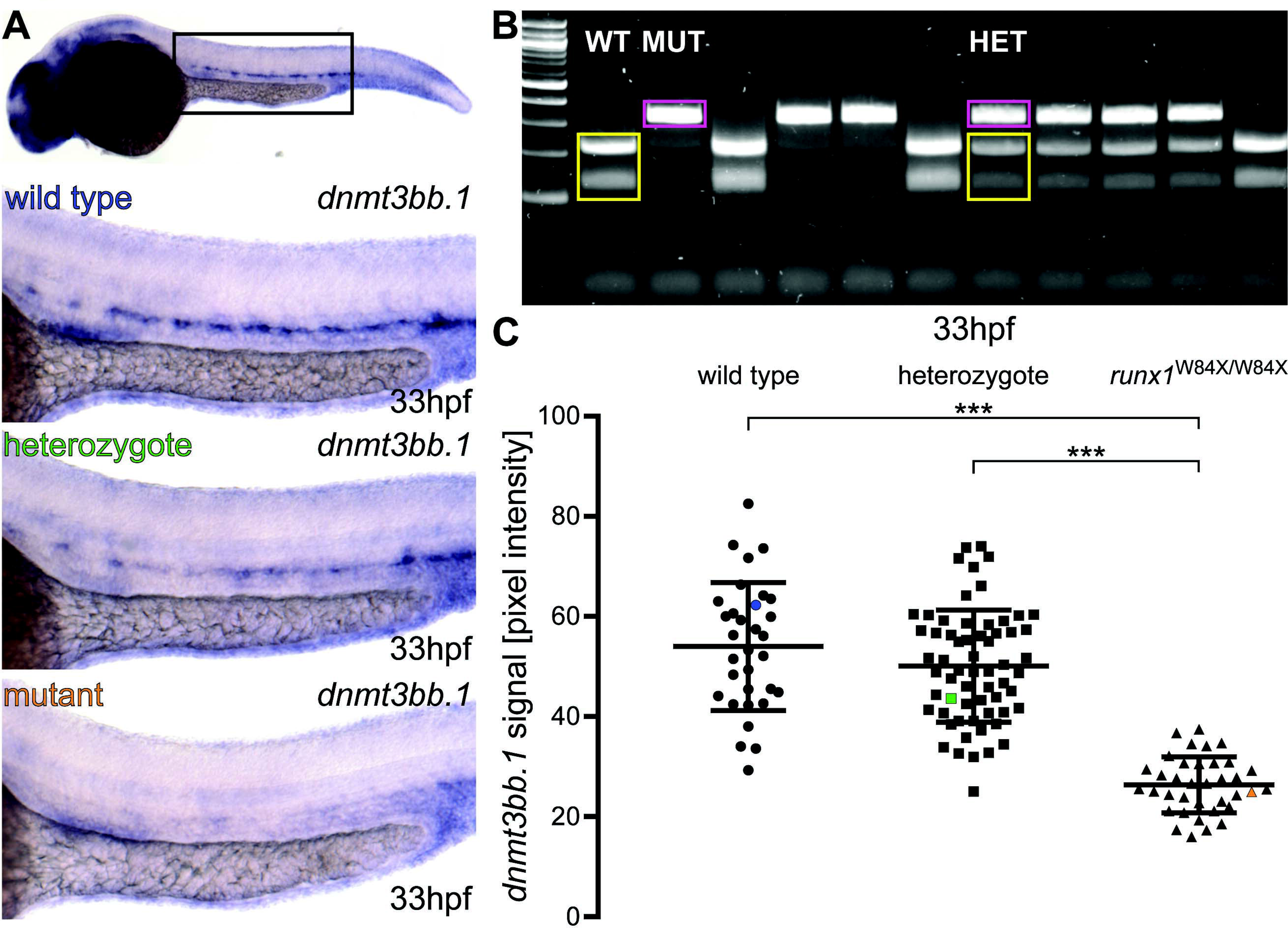
*runx1* mutants have significantly reduced levels of *dnmt3bb.1* mRNA detected by ISH. A) Representative images of ISH for *dnmt3bb.1* in 33hpf wild type (blue), *runx1*^+/W84X^ (green) and *runx1*^W84X/W84X^ (orange) embryos, showing the expression in the dorsal aorta. The top panel indicates the location of the zoomed regions in a wild type embryo. B) 2% agarose gel showing representative genotypes of wild type (WT), heterozygous (HET) and mutant (MUT) *runx1* embryos, distinguished by RFLP. Yellow: wild type 214bp + 124bp bands, pink: 338bp mutant band. First lane from the left: 100bp DNA ladder. C) Pixel intensity values of *dnmt3bb.1* mRNA in *runx1*^W84X/W84X^embryos (n=36) are significantly decreased compared to wild types (n=32) and heterozygotes (n=62) (ANOVA, p<0.001). The coefficients of variation are 24%, 22% and 21% for wild type, heterozygote and mutant groups, respectively. Blue, green and orange data point correspond to the example images from panel A. The bars represent mean ± s.d. \*\*\**p*<0.001 (Games-Howell post-hoc test).

We applied the same analysis method to the images of 16 *gpr65* morphants and 16 control (uninjected) siblings probed at 29hpf for *runx1* mRNA, previously shown to be negatively regulated by *gpr65* (Gao et al., 2016). We found a significant increase of approximately 25% in pixel intensity levels of the *runx1* probe staining in *gpr65* morphants, compared to non-injected controls (μ_wt_=29, μ_gpr65_=37.2; *t*=2.38, d.f.=30; *p*<0.05) (Fig. 3). The values of the wild type embryos and the morphants showed dispersion with the coefficients of variation of 34% and 26%, respectively.

**Figure 3.**
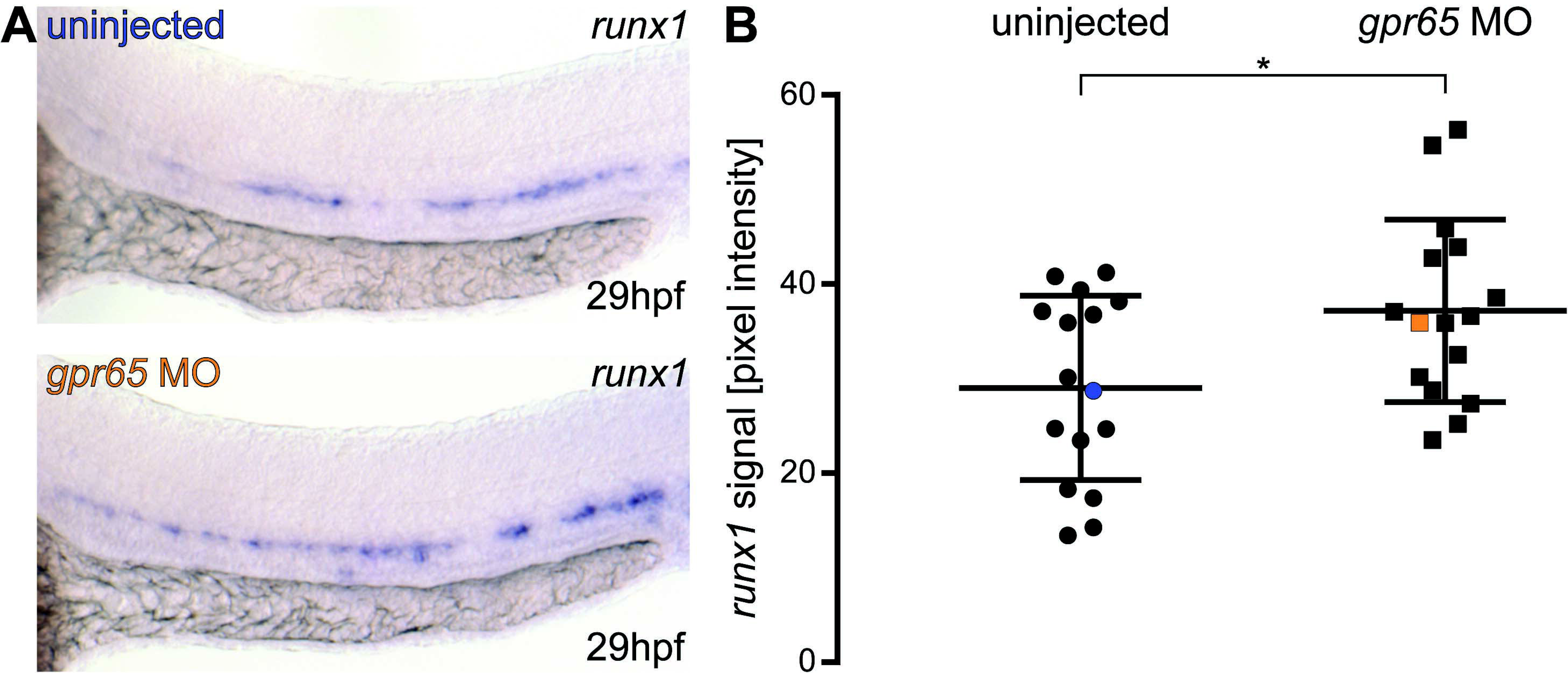
*Gpr65* morphants have significantly increased levels of *runx1* mRNA detected by ISH. A) Representative images of ISH for *runx1* in 29hpf wild type (blue) and *gpr65* MO-injected (orange) embryos, showing the expression in the dorsal aorta. B) Pixel intensity values of *runx1* mRNA in in *gpr65* MO-injected embryos (n=16) are significantly higher than in uninjected control siblings (n=16). The coefficients of variation are 34% and 26% for wild type and morphant groups, respectively. Blue and orange data point correspond to the example images from panel A. The bars represent mean ± s.d. \**p*<0.05 (t test). The power of the *t* test to detect the difference at 0.05 level was 63%.

### Digital quantification of runx1 ISH signal in the dorsal aorta reveals a discrepancy between the phenotype in lmo4a^Δ25-29/Δ25-29^ mutants and previously described lmo4a morphants

We applied the ISH quantification and genotyping methods to analyse the phenotype of *lmo4a* mutants. MO-mediated knockdown of *lmo4a* had previously been shown to result in reduced expression of *runx1* in the dorsal aorta at 30hpf (Meier et al., 2006). We used TALENs to generate a 5bp deletion in the coding region of the *lmo4a* gene *(lmo4a^Δ25-29^),* which is predicted to be a null (Fig.4A). We then incrossed *lmo4a* heterozygous parents and quantified the expression of *runx1* in 28hpf embryos prior to genotyping them with 100% efficiency using RFLP (Fig. 4B-C). Surprisingly, when comparing *runx1* expression in these mutants, we found no significant differences in *runx1* pixel intensity at 28hpf among wild type, heterozygous and homozygous siblings (μ_wt_=30.3, μ_het_=32.8, μ_mut_=34.6; *F*=0.509, d.f.=2, 62; *p*>0.6) (Fig. 4D). The coefficients of variation were 41%, 38% and 37%, respectively. We also compared *runx1* mRNA levels in single wild type and *lmo4a* mutant embryos by qRT-PCR. These experiments revealed a small (27%) yet significant decrease in *runx1* mRNA levels in *lmo4a* mutants compared to wild type (ΔCt_wt_=12.44, ΔCt_mut_=12.83; *t*=2.427, d.f.=34; *p*<0.05) (Fig. 4E). This result may indicate that in certain cases digital quantification is unable to detect small differences in *mRNA* expression levels; yet we cannot rule out the possibility that *runx1* from other sources contributes to the decrease observed in the qRT-PCR.

**Figure 4.**
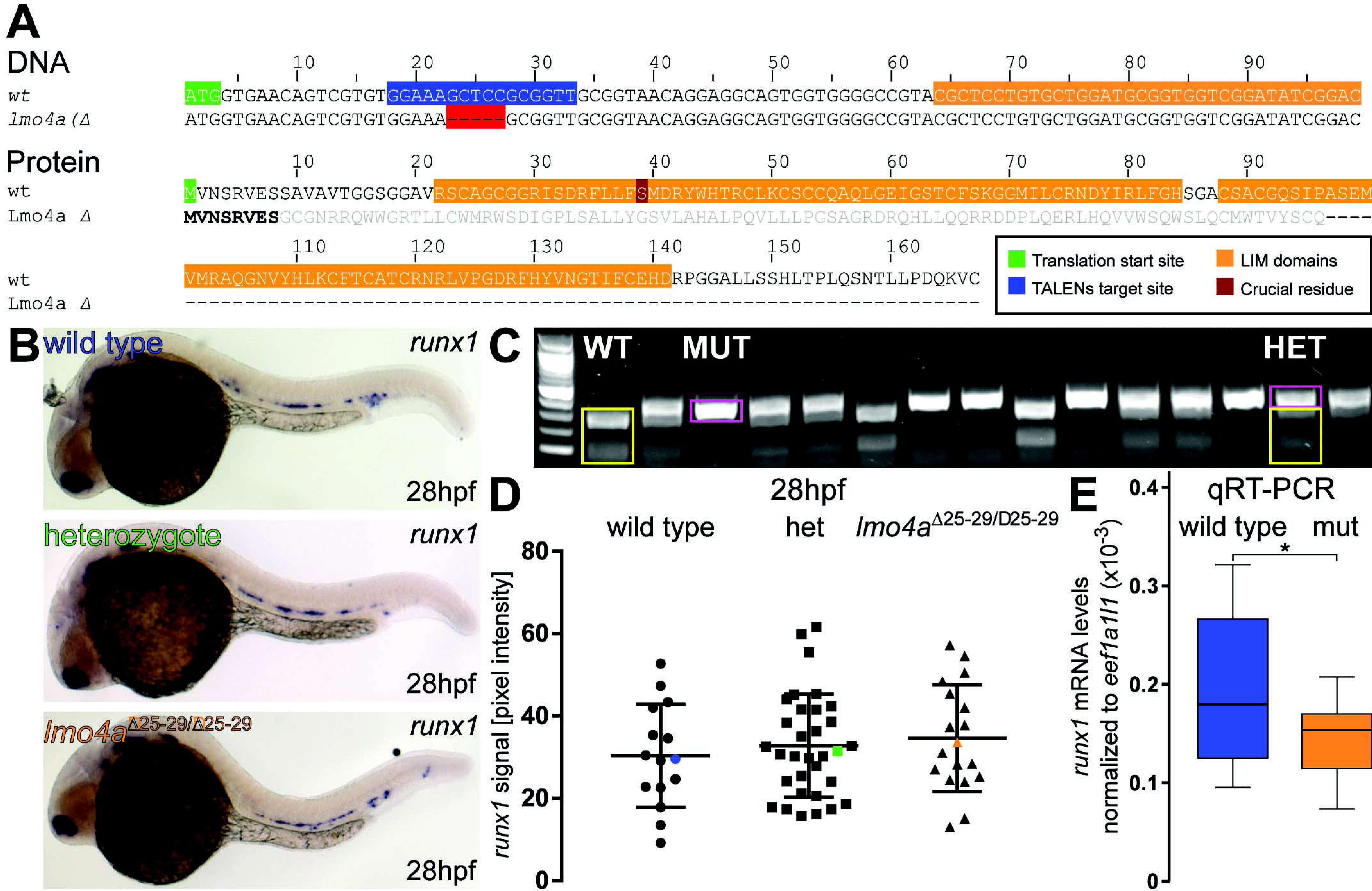
*runx1* levels detected by ISH are not affected in *lmo4a* mutants. A) TALENs were designed to a region (blue) ∽20bp downstream of the *lmo4a* translation start site (green). Isolated mutant alleles carry 5bp deletions (Δ25-29) (red gaps) upstream of the conserved LIM domains (orange), resulting in a frameshift after S8. The resulting mutant protein is predicted to lack the LIM domains, including the crucial S39 (purple). B) Representative images of ISH for *runx1* in 28hpf wild type (blue), heterozygous (green) and lmo4a^Δ25–29/Δ25–29^ (orange) embryos, showing the expression in the dorsal aorta. C) 2% agarose gel showing representative genotypes of wild type (WT), heterozygous (HET) and mutant (MUT) *lmo4a* embryos, distinguished by RFLP. Yellow: wild type 258bp + 87bp bands, pink: 340bp mutant band. First lane from the left: 100bp DNA ladder. D) Quantification of the *runx1* mRNA signal, detected by ISH, from 28hpf wild type (n=15), heterozygous *lmo4a^+/−^* (het) (n=34) and lmo4a^Δ25–29/Δ25–29^ mutant (n=18) embryos from one clutch shows no significant difference in *runx1* pixel intensity among the different genotypes (*t* test, *p*>0.6). The coefficients of variation are 41%, 38% and 37% for wild type, heterozygote and mutant groups, respectively. Blue, green and orange data point correspond to the example images from panel B. The bars represent mean ± s.d. E) Boxplots displaying normalized *runx1* mRNA levels (2^-ΔCt^) in single wild type (blue; n=12) and lmo4a^Δ25–29/Δ25–29^ (mut, orange; n=12) embryos, measured by qRT-PCR, showing decreased levels of *runx1* in the mutants compared to wild type. \**p*<0.05 (*t* test).

## DISCUSSION

The HotSHOT method of genomic DNA isolation, originally designed for mouse ear notch samples (Truett et al., 2000), offers a fast and cost-effective way to genotype animals. While it has been used to isolate genomic DNA from paraformaldehyde-fixed zebrafish samples before (Cooney et al., 2013; Meeker et al., 2007), its efficiency in this setting had not been reported. In consequence, the method has not been widely adopted in the community and many research groups rely on time-consuming and more expensive DNA extraction methods involving proteinase K treatment (Gore et al., 2016) or commercially available kits (Bresciani et al., 2014; Sood et al., 2013). Here we report that PCR-quality DNA can be rapidly extracted from 95-100% of fixed zebrafish embryos aged from 14hpf to 5dpf with an optimised HotSHOT protocol after ISH. This DNA can subsequently be used to genotype the samples with simple, inexpensive PCR followed by a digest to detect restriction enzyme sites disrupted by the mutation, as done previously for fresh tissue (Hruscha et al., 2013). To facilitate this, mutations can be designed to target restriction enzyme recognition sites. In fact, an online TALEN and CRISPR/Cas9 design tool Mojo Hand (Neff et al., 2012) readily provides restriction enzyme sites targeted by a desired mutation. While newly emerging alternatives to standard PCR and RFLP may provide a higher speed of genotyping (D’Agostino et al., 2016; Lee et al., 2016), the protocol described here for genotyping after ISH is attractive due to its high efficiency and demonstrated robustness in our hands.

Genotyping zebrafish embryos after ISH is important because it allows processing of mutant and wild type embryos in one batch, therefore limiting the technical variation between samples. In addition, expression levels of the target gene can be assessed in a non-biased way, because the embryos can be distinguished by their genotype only after phenotypic assessment. This is a powerful way to control for unconscious bias, a serious issue in *in vivo* animal research (Macleod et al., 2015). Assessing mRNA levels in post-ISH embryos has been performed visually, either by scoring the phenotypes into discrete groups (Blaser et al., 2017; Gao et al., 2016; Genthe and Clements, 2017) or by cell counting (Espín-Palazón et al., 2014). However, these approaches are prone to subjectivity and poor reproducibility. Furthermore, visual scoring can be difficult to carry out and interpret due to expression level differences between individuals of the same genotype. Indeed, we show here that pixel intensities of the ISH signal in wild type embryos probed for the transcription factor *runx1* show high dispersion in wild type embryos with over 25% and up to 40% coefficient of variation (Figs 3B, 4D). These results indicate that the interpretation of phenotypes based purely on expected Mendelian distribution from heterozygous incrosses might be misleading. As we demonstrate, visual scoring of embryos from a heterozygous *runx1^+/W84X^* incross into ‘high’, ‘medium’ and ‘low’ groups based on *dnmt3bb.1* expression levels gives a phenotypic Mendelian distribution of 1:2:1, which could suggest a haploinsufficiency effect on the regulation of *dnmt3bb.1* expression. Genotyping of these embryos revealed that the vast majority of ‘low’-expressing ones were indeed genetically homozygous mutant. However,‘high’- and ‘medium’-expressing embryos were distributed similarly across wild type and heterozygous fish, disproving the haploinsufficiency hypothesis. As a possible explanation for this discrepancy, we found that the signal intensity values in all three genotypes were highly dispersed, with coefficients of variation over 20%. Therefore, each ISH experiment done on embryos from a heterozygous incross should be followed by genotyping to avoid misleading conclusions due to the variability of the ISH signal intensities in embryos of the same genotype.

We would argue that the use of digital image analysis on ISH-probed samples is critical for objective, statistical demonstration of changes in expression levels. Here we describe an imaging protocol based on previous studies (Fan et al., 2015; Wen et al., 2017) to measure gene expression intensity in the trunk region of 1-2 day old zebrafish embryos. We show that the average ISH staining intensity for *dnmt3bb.1* mRNA is significantly decreased in *runx1* mutants compared to wild type siblings, in agreement with previously reported qRT-PCR quantitation of *dnmt3bb.1* levels in whole embryos (Gore et al., 2016). Thus, our method is robust and should be adopted instead of less reliable visual scoring methods. It could also be used as an alternative to qRT-PCR experiments where these require larger numbers of animals and are prone to errors due to a limited number of highly reliable internal controls (Xu et al., 2016). Our quantification method addresses all of these limitations. Furthermore, it presents a way to measure changes in expression levels in a very tissue-specific manner, which is useful in the case of genes with multiple developmental roles. We believe it will be particularly helpful for studying other genes with expression patterns that are spatially restricted, such as *gata2b,* a haematopoietic gene expressed in the ventral wall of the dorsal aorta (Butko et al., 2015). An alternative modification to our method could involve subtracting the average intensity of another stained region from the intensity of the selected ROI – for instance, using the *runx1* or *dnmt3bb.1* signal in the head as an internal control. However, expression levels vary widely between different tissues and there is a risk of saturating the signal, reducing the dynamic range used for the comparisons. Therefore, we believe that using an unstained region with the same area as a background measurement provides a more reliable way to quantify the ISH signal in each embryo. When measuring *runx1* staining intensity in the dorsal aorta, we chose an unstained region dorsal to the notochord as background. We found the pixel intensity values of this region remarkably stable across experiments with a 10% coefficient of variation. In extreme cases, the background pixel intensity value of this area is so high that subtraction from the signal value produces a negative number. However, these are very rare instances (0.4% of *runx1*-probed embryos per experiment) and thus this limitation is unlikely to influence the overall outcome.

We also propose a way to quantitatively represent variation in gene expression levels without relying on subjective and biased scoring. For instance, we could replicate the previously reported increase in *runx1* expression in *gpr65* morphants (Gao et al., 2016) with our method, but we represented it in a more objective quantitative way, importantly allowing statistical analysis. In fact, we achieved 63% power to detect a 25% increase in pixel intensity at the *p*<0.05 level for sample sizes as small as 16 for each condition. This method of analysis also allows precise calculations of required sample sizes to achieve a given power. In the presented example, 90% power would require 31 MO-injected embryos and 31 uninjected controls. Such calculations are essential in animal research (Dell et al., 2002), but they are notoriously not included (Macleod et al., 2015). In fact, recently published updated guidelines for the use of MOs in zebrafish encourage statistical analysis of phenotypes and advocate the use of blinded assessment (Stainier et al., 2017) and both points can be addressed with our method. In addition, there is scope to automate the phenotypical analysis using this method. We have generated a batch conversion Fiji macro that converts all images in a given directory (see Supplementary Detailed Protocol) ready for manual ROI selection. Future optimisations could involve automation of intensity measurements of the ISH images (Chen et al., 2011).

We have further applied the described method to analyse the molecular phenotype of an unpublished *lmo4a* mutant. A previous report had shown decreased *runx1* expression in *lmo4a* morphants (Meier et al., 2006). By contrast, we found no difference in *runx1* pixel intensity levels between wild types, heterozygotes and *lmo4a* homozygous mutants. These findings were not supported by our single embryo qRT-PCR experiments, where we found a 27% decrease in *runx1* mRNA levels in *lmo4a* mutants. Because the qRT-PCR experiment was performed on whole embryos, it is likely that the resulting decrease is due to the expression of *runx1* in other parts of the embryo (nasal placodes and neurons, for example). While we cannot rule out that digital quantification might not be sensitive enough for detecting small changes in expression levels in all cases, it is important to note that the strength of image quantification is its specificity to the area and tissue of interest. Taken together, our results highlight that when assessing small differences in mRNA expression levels, the results should be cross-validated using methods that enrich for cells or tissues of interest e.g. FACS sorting based on marker expression followed by qPCR.

## METHODS

### Maintenance of zebrafish and morpholino oligonucleotide injections

All animal experiments were approved by the local ethics committee. Wild type and *runx1*^W84X^ (Jin et al., 2009) and *lmo4a^Δ25-29^* mutant zebrafish *(Danio rerio* Hamilton) were maintained and bred according to standard procedures (Westerfield, 2007). Embryos were collected by natural mating of 4-18-month old adults and staged according to morphological features (Kimmel et al., 1995) corresponding to respective age in hours or days post fertilisation (hpf or dpf, respectively). For *gpr65* knockdown, wild type one-cell stage embryos were injected with 4ng of GPR65_SP MO (Gao et al., 2016).

### Generation of lmo4a mutants

For TALENs design, Mojo Hand software was used (http://www.talendesign.org)(Neff et al., 2012). The identified target site was GGAAAGCTCCGCGGTT. The RVD-containing repeats were assembled using the Golden Gate approach in pTAL3-DDD and pTAL3-RRR vectors (Cermak et al., 2011). The resulting DNA templates were verified by sequencing and linearised with NotI enzyme. The mRNAs were transcribed from 1μg linearised template with SP6 mMessage mMachine^®^ kit (Ambion) and purified with RNeasy^®^ Micro Kit (QIAGEN).

One-cell zebrafish embryos were injected with 100pg left-arm + 100pg right-arm TALEN mRNAs. Germline mutations in the founders (lmo4a^+/−^) were identified using restriction fragment length polymorphism (RFLP)(Bedell et al., 2012) (see primer sequences below) with SacII enzyme on the genomic DNA extracted from their offspring. Mutations were identified by the presence of an undigested PCR product on a 2% agarose gel. The undigested mutant fragments were purified from the gel, cloned into pGEM^®^-T Easy vector (Promega) and sequenced. The *lmo4a^+/−^* founders were outcrossed to generate heterozygous carriers, identified with fin-clipping and genotyping at 3dpf as described previously (Wilkinson et al., 2013).

### Whole-mount in situ hybridization

ISH was carried out according to the standard lab protocol (Jowett and Yan, 1996) using digoxygenin-labelled *dnmt3bb.1* (Gore et al., 2016) and *runx1* (Kalev-Zylinska et al., 2002) probes. Post hybridization, the embryos were bleached in 5% formamide/0.5% SSC/10% H_2_O_2_ (Monteiro et al., 2011) and imaged in 100% glycerol with QImaging MicroPublisher 5.0 RTV Camera and Q-Capture Pro 7^™^ software (version 7.0.3), using the same exposure, magnification and illumination settings for each embryo.

### DNA extraction and genotyping

Genomic DNA was isolated from 4% paraformaldehyde (PFA)-fixed embryos using the original HotSHOT protocol (Truett et al., 2000) (Fig. 1a). Briefly, 40-75 μl of lysis buffer (25 mM NaOH, 0.2 mM EDTA) was added directly to a PCR tube with a freshly-imaged embryo in <5 μl 100% glycerol. To test the efficiency of the DNA extraction, embryos were suspended in the buffer and incubated at 95°C for 5-120 minutes, then cooled to 4°C, after which an equal volume of neutralisation buffer (40-75 μl 40 mM Tris-HCl) was added (see detailed Supplementary protocol). Genomic regions containing the mutated sites in the *runx1* locus were amplified with JumpStart^™^ REDTaq^®^ ReadyMix^™^ PCR or with Phire^™^ Green HotStart II PCR Master Mix according to manufacturer protocols, using 5 μl of DNA lysate in a 20 μl reaction volume and the following primer sequences: 5’-GCTCTGGTGGGCAAACTG-3’ and 5’-CATGTGTTTGGACTGTGGGG-3’ for *runx1;* and 5’-ACTTTGCCTCTGGATCTGCT-3’ and 5’-TCTACATCAGTCCCGCCAAA-3’ for *lmo4a.* The presence of *runx1* and *lmo4a* mutations were verified by restriction fragment length polymorphism (RFLP)(Hruscha et al., 2013) with HaeII or SacII enzymes respectively on a 2% agarose gel, using 100bp DNA Ladder as a reference. The sequences of additional primers used were: 5’-GCACCACAGTGGACATTGAT-3’ and 5’-GTTGTAGAGGGCCAGCACTT-3’ for the *tgfβ3* locus; 5’-CATTAATGCGAGGGATACGG-3’ and 5’-AAAAGAGCCACGGTAGGTGA-3’ for the *tgfβr2* locus; and 5’-TGGCTAAGTGACCGTCAGAG-3’ and 5’-TGAAACAAAACGCAGACGAC-3’ for the *gata2a* locus.

### Digital image analysis

Using Fiji software (Schindelin et al., 2012), the images were inverted to negative and converted to 8-bit grayscale. A Region of Interest (ROI) containing the ISH expression signal in the dorsal aorta along the yolk sac extension was drawn manually for each embryo. Then a second ROI with the same shape and area was created in a region of the embryo that has a uniform intensity and does not contain any ISH staining. In this particular instance, this region was placed just above the notochord (Fig. 1A). This area was used to define the background. A value for each region was then determined by measuring the average pixel intensity. After subtracting the value of the background region from the value of the stained region, the pixel intensity of the ISH signal was assigned to each embryo (Fig. 1A and detailed Supplementary protocol). The graphs presenting individual data points, means and ±s.d. were plotted using GraphPad Prism 7.

### Quantitative RT-PCR

Total RNA was extracted from single wt and *lmo4a^-/-^* embryos using TRIzol^™^ Reagent (Invitrogen) and Direct-zol^™^ RNA MiniPrep kit (Zymo Research). cDNA synthesis was performed with the Superscript IV RT kit (Invitrogen) and diluted ¼ in H_2_O. For qRT-PCR, we used 3μl of diluted cDNA per sample (in triplicate) in 20 μl reactions containing the Fast SYBR^™^ Green Master Mix (Thermo Fisher Scientific) and a pair of the following primers: *runx1*: 5’-TTCACAAACCCTCCTCAAG-3’ and 5’-CTGCTCAGAGAAAGCTAACG-3’; *eef1a1/1:* 5’-GAGAAGTTCGAGAAGGAAGC-3’ and 5’-CGTAGTATTTGCTGGTCTCG-3’. The reactions were run on a 7500 Fast Real-Time PCR System (Applied Biosystems). The relative mRNA levels in each sample were calculated by subtracting the mean of Ct values for the housekeeping gene *eef1a1/1* from the average Ct values for *runx1.* These values (ΔCt) were then converted to a ratio relative to *eef1a1/1* with the formula 2^-ΔCt^. The graphs were plotted using IBM^®^ SPSS^®^ Statistics (version 22) software.

### Statistical analysis

The numbers of embryos scored as ‘high’, ‘medium’ or ‘low’ were tested for equal distribution among wild type, heterozygous and mutant genotypes with contingency Chi-squared tests, applying Continuity Correction for 2×2 tables. For digitally analysed images, the pixel intensity values were assessed for normal distribution with Q-Q plots and transformed with sqrt function if necessary. Mean values (μ) of each experimental group were analysed with 2-tailed independent-samples t-test (for *gpr65* MO experiment) or with ANOVA (for mutant experiments) with 95% confidence levels, testing for the equality of variances with a Levene’s test and applying the Welch correction when necessary. For ANOVA, differences between each two groups were assessed with either Tukey’s post-hoc test (for equal variances) or with Games-Howell test (for unequal variances). The degree of variability in each sample was assessed by calculating the coefficients of variation, defined as 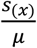 with *s_(x)_* being the standard deviation. The post-hoc power of the tests and required sample sizes were determined with G^*^Power software (version 3.0.10)(Faul et al., 2007).

The ΔCt values from qRT-PCR experiments were assessed for normal distribution with a Q-Q plot and analysed with 2-tailed independent-samples t-test with 95% confidence levels, testing for the equality of variances with a Levene’s test. For all analyses the IBM^®^ SPSS^®^ Statistics (version 22) package was used.

## ACKNOWLEDGEMENTS

We thank Dr Dominic Waithe from the Wolfson Imaging Centre for advice on developing the image quantification method. We thank Rossella Rispoli for help with generating the batch conversion Fiji macro. We thank staff members of Biomedical Services at the John Radcliffe Hospital for monitoring and feeding zebrafish.

## DATA AVAILABILITY

All data generated or analysed during this study are included in this published article (and its Supplementary Information files).

